# The Anaeramoeba symbiosome: a single contiguous organelle that doubles the cell’s membrane surface

**DOI:** 10.64898/2026.04.10.717692

**Authors:** Jon Jerlström-Hultqvist

**Affiliations:** Department of Cell and Molecular Biology, BMC, Box 596, Uppsala Universitet, Sweden, SE-751 24 Uppsala, Sweden

**Keywords:** *Anaeramoeba*, symbiont, organelle, syntrophy, symbiosis, FIB-SEM, *Desulfobacter*, symbiosome, sulfate-reducing, hydrogenosomes

## Abstract

Anaerobic protists across diverse lineages have independently evolved intimate spatial associations between their hydrogen-producing mitochondrion-related organelles and prokaryotic symbionts, yet the cellular structures mediating these syntrophic partnerships remain poorly characterized. Anaeramoebae — a recently described phylum of anaerobic amoeboflagellates — have evolved a particularly elaborate solution: the symbiosome, a membrane organelle that houses sulfate-reducing *Desulfobacter* sp. symbionts alongside host hydrogenosomes and maintains direct connections to the extracellular environment. Previous FIB-SEM work using aldehyde-based fixation established the symbiosome as a dynamic structure but left critical architectural questions unresolved, including whether symbionts occupy separate compartments and how extensive the connections to the cell exterior truly are. Here, we use high-pressure freezing with optimized cultivation to achieve markedly improved membrane preservation in *Anaeramoeba flamelloides*. We show that the symbiosome is a single, fully interconnected compartment enclosing all *Desulfobacter* sp. symbionts, spanning up to 15% of total cell volume with a membrane surface area matching that of the plasma membrane. The number of symbiosome-to-surface connections is an order of magnitude higher than previously documented — 12 and 29 pores in two cells, compared with three in an earlier published volume — likely reflecting the metabolic requirement for extracellular sulfate access by the symbionts. These findings establish the *Anaeramoeba* symbiosome as one of the largest known membrane organelles in a single-celled eukaryote, with an architecture shaped by the demands of syntrophic exchange.

## Main text

Anaerobic protists are ecologically important predators of prokaryotes and major drivers of nutrient cycling in oxygen-depleted environments^1^. Many have independently evolved symbiotic partnerships with bacteria or archaea^2^, sustained by the metabolic output of modified mitochondria^3^ — mitochondrion-related organelles (MROs) — that produce hydrogen, acetate, and other compounds as fermentation end-products. A defining feature of these systems is the close spatial association between MROs and their symbiotic partners, enabling direct interspecies transfer of H_2_ and other metabolites in a process called syntrophy^4,5^. Hydrogen removal is mutually beneficial: it relieves product inhibition of host fermentation while providing electron donors for symbiont metabolism. Superficially similar arrangements of hydrogenosomes and symbionts have been documented in anaerobic ciliates^6,7^, the heterolobosean *Psalteriomonas lanterna*^8^, and obligately symbiotic parabasalids in the termite gut^9,10^, where prokaryotic partners — often methanogens or sulfate-reducing bacteria — are housed in shallow host-derived membrane invaginations with hydrogenosomes in close proximity. Yet despite this recurring theme, the cellular structures that establish and maintain these associations remain poorly characterized. Anaeramoebae — a recently described phylum of anaerobic amoeboflagellates^11^ — have evolved a particularly elaborate solution: the symbiosome, a membrane organelle that co-localizes sulfate-reducing *Desulfobacter* sp. symbionts with host hydrogenosomes and maintains direct connections to the extracellular environment, physically integrating the syntrophic partnership within the architecture of a single cell^12,13^. Symbiosomes have evolved on numerous occasions across the tree of life, and their origins are associated with profound changes to host cell biology^14^. In *Anaeramoeba*, the notable expansion of membrane-trafficking gene families in the common ancestor suggests that the symbiosome is a highly derived structure that enables selective management of captured symbionts^13^. Indeed, recently described *Anaeramoeba* strains display additional, distinct modes of prokaryotic symbiosis, underscoring the diversity of symbiont–host interfaces within this single phylum^15^.

Preparing marine anaerobic cells for electron microscopy poses particular challenges for preserving membranous structures with biological fidelity. High-pressure freezing enables rapid vitrification, but the choice of cryo-protectant is critical for achieving preservation without freezing damage. Previous FIB-SEM work on *Anaeramoeba* relied on aldehyde-based chemical fixation, which resulted in compromised membrane preservation and potential fixation artefacts that limited the ability to trace fine membrane connections^13^. Those experiments documented only three symbiosome-to-surface connections and, although a majority of symbionts resided in a single compartment, several apparently disconnected subcompartments were observed. Here, we optimize the growth and fixation conditions for *Anaeramoeba flamelloides* BUSSELTON2, using high-pressure freezing with 20% BSA as cryo-protectant, to overcome these fixation-related limitations.

We imaged and segmented two nearly complete *A. flamelloides* BUSSELTON2 cells by FIB-SEM (volumes 9235_1 and 9235_2). Cells were grown directly on sapphire discs under biologically generated anaerobic conditions, yielding well-vitrified specimens without evidence of freeze damage. The resulting membrane preservation was markedly improved: the plasma membrane appeared intact and smooth, without the jaggedness observed in the previous volume^13^. Both cells displayed expected *Anaeramoeba* features, including a droplet-shaped nucleus with prominent pores, a ventral microtubule-organising centre, and an extensive endomembrane system (Figure S2, Table S1).

The symbiosome was the dominant intracellular structure in both cells. Cell one contained 160 *Desulfobacter* sp. symbionts within a symbiosome occupying 15.31% of total cell volume; cell two contained 95 symbionts in a symbiosome occupying 11.08% (Table S1). The symbionts are elongated rods (Figure S1), seven and four of which (in cells one and two, respectively) were in various stages of cell division. The bacterial symbionts accounted for 6.95% and 5.11% of cell volume in cells one and two, respectively. Hydrogenosomes were numerous — 374 organelles (6.31% of cell volume) in cell one and 307 (6.1%) in cell two — and tightly positioned along the symbiosome membrane, occupying the space between symbiosome projections (Figure 1). In cell one, the symbiosome envelops the dorsally projecting nucleus, whereas in cell two, the two structures are spatially separated. Strikingly, despite the symbiosome occupying 15.31% and 11.08% of total cell volume, its membrane surface area (2.32% and 1.61% of cell volume) is roughly equivalent to that of the plasma membrane (2.21% and 1.69%), effectively doubling the cell’s total exposed membrane surface.

**Figure 1:**
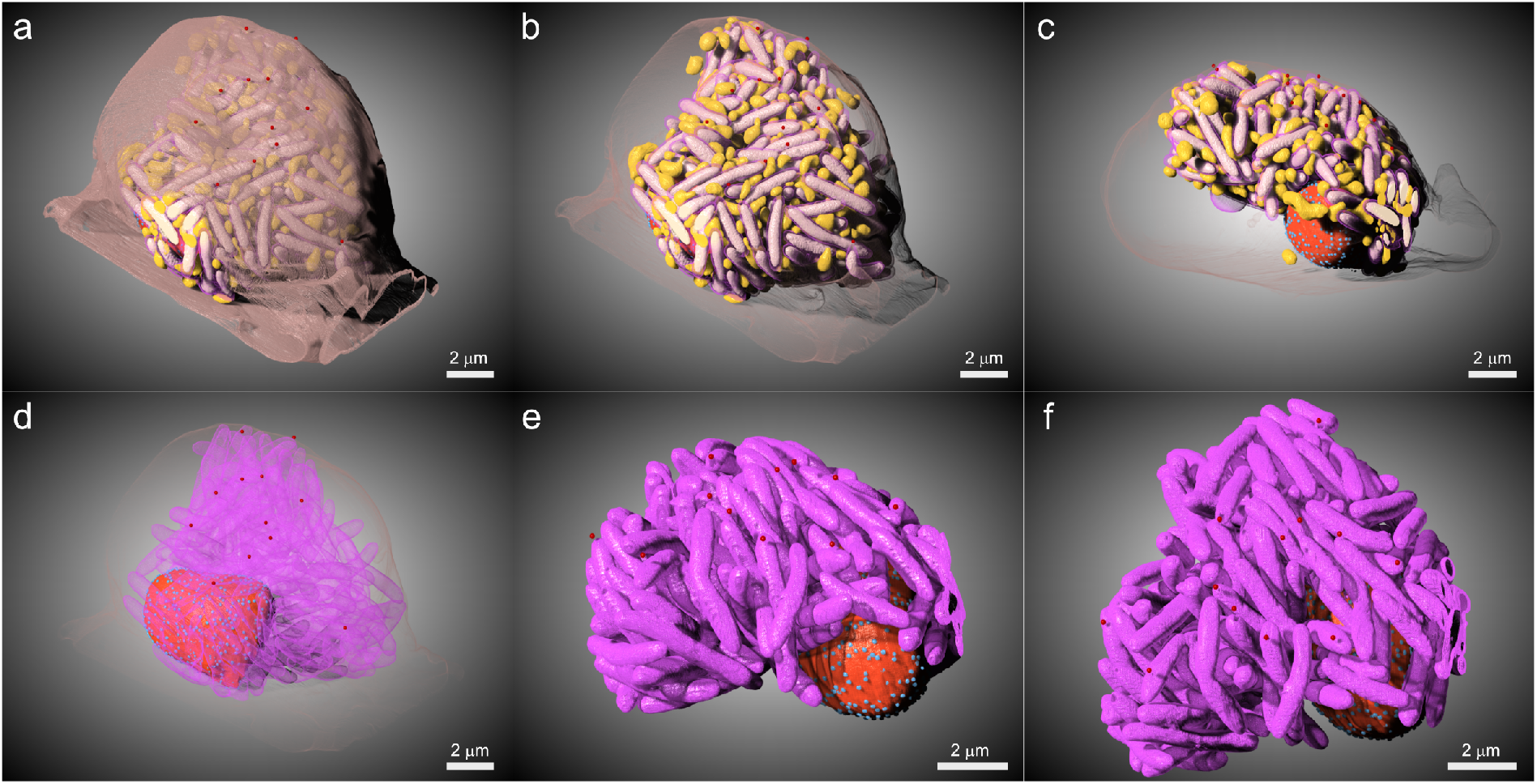
High-pressure frozen cell of *A. flamelloides* BUSSELTON2 with well-preserved membrane-structures. **a-f**, Renderings of selected segmented organelles/cell structures of *A. flamelloides* BUSSELTON2 volume 9235_1, showing (symbiont – white, hydrogenosome – yellow, symbiosome membrane – purple, nucleus – red, nuclear pores – turquoise, plasma membrane – light brown, symbiosome-to-surface openings – dark red). **a**, symbiont, hydrogenosome, symbiosome membrane, nucleus, nuclear pores, plasma membrane, symbiosome-to-surface openings. **b**, as in **a**, but with plasma membrane shaded. **c**, same as in **b**, but the cell is facing sideways. **d**, symbiosome membrane shaded, nucleus, nuclear pores, symbiosome-to-surface openings, and plasma membrane shaded. **e**, symbiosome, nucleus, nuclear pores, and symbiosome-to-surface openings with the cell facing sideways. **f**, as in **e**, but the cell is being viewed from the top. Scale bar 2 µm.

Critically, in both volumes, the symbiosome was fully interconnected, with all symbionts residing in a single contiguous compartment — resolving the question of whether the disconnected subcompartments observed previously^13^ reflected genuine cellular architecture or fixation artefacts. The improved preservation also revealed far more extensive connections between the symbiosome and the cell exterior than previously appreciated. Where the earlier volume documented only three pore-like openings^13^, we identified 12 (Figure 2AB) and 29 (Figure 2CD) symbiosome-to-surface connections in cells one and two, respectively (Figure S3). The abundance of these connections likely reflects a metabolic constraint: the sulfate-reducing *Desulfobacter* sp. symbionts require access to extracellular sulfate, paralleling the situation in *Desulfovibrio* symbionts of termite-gut *Trichonympha*, where surface connections to the symbiosome are similarly maintained^9,10^. Whether these pores are relatively static or continuously forming and closing remains unclear, but their number and distribution suggest active regulation. Our finding that the symbiosome is a single, fully interconnected membrane compartment is consistent with previous pulsed labelling experiments showing that fluorescently labelled WGA rapidly stains the entire symbiont population^13^.

**Figure 2:**
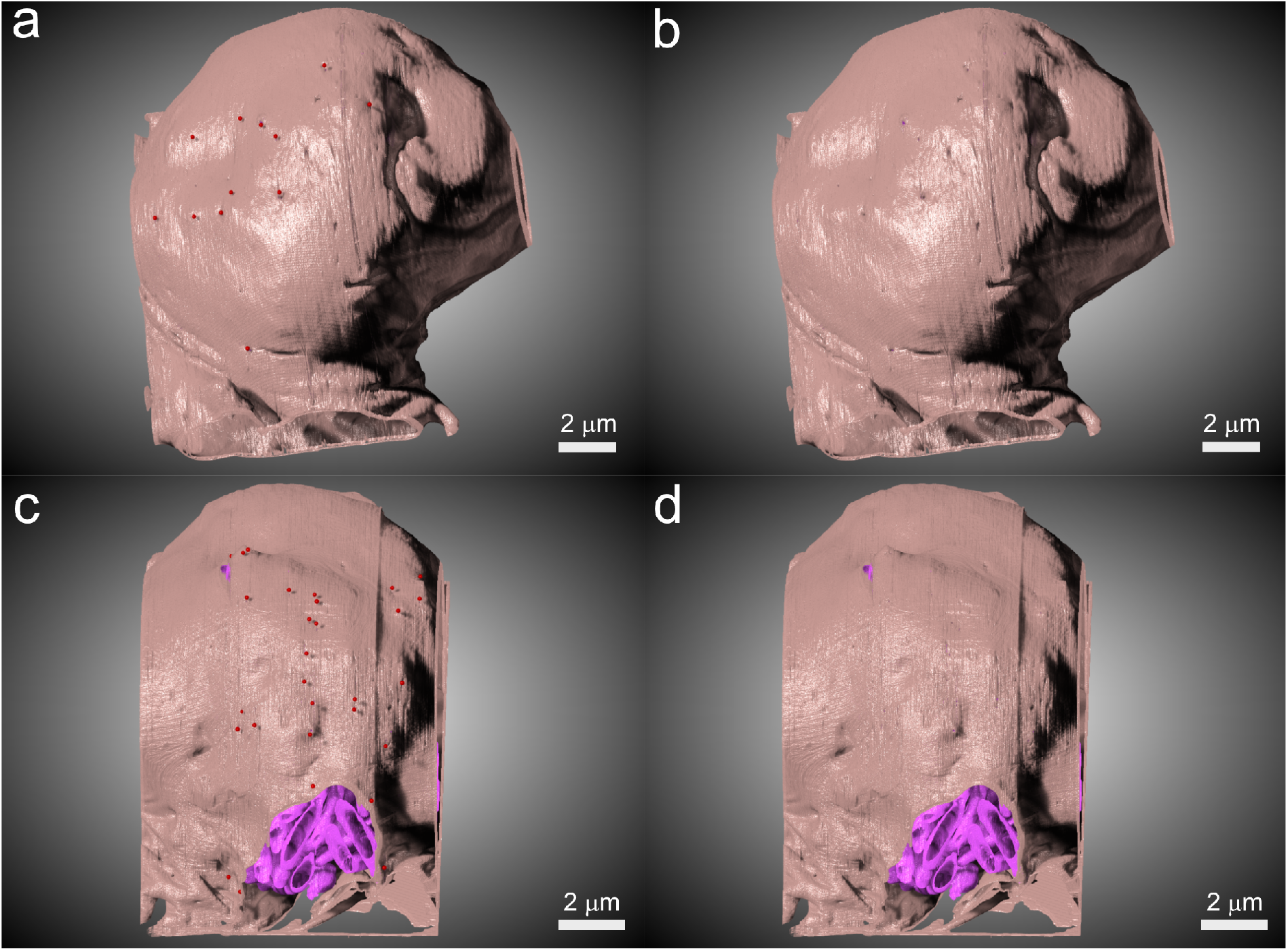
Symbiont contact distribution on the *A. flamelloides* BUSSELTON2 cell membrane. **a-b**, Cell 9235_1: **a**, plasma membrane with symbiosome-to-surface openings (red spheres), **b**, plasma membrane alone. **c-d**, Cell 9235_2: **c**, plasma membrane with symbiosome visible and symbiosome-to-surface openings indicated, **d**, plasma membrane with symbiosome visible. symbiosome membrane – purple, plasma membrane – light brown, symbiosome-to-surface openings – dark red. Scale bar 2 µm.

Together, these data establish the *Anaeramoeba* symbiosome as a massive organellar membrane system — spanning up to 15% of total cell volume with a surface area matching the plasma membrane and connected to the extracellular environment through numerous pores — that physically couples hydrogenosomal hydrogen production with symbiont sulfate reduction within a single eukaryotic cell.

## Materials and methods

### Cell cultivation

*A. flamelloides* BUSSELTON2 cells were routinely passaged and grown as described in Jerlström-Hultqvist et al^13^. Briefly, newly split cells (50% media replacement) were grown in Artificial Seawater + 3% Luria Bertani broth for 2 days into late exponential phase in a 75 cm^2^ Nunc™ EasYFlask™ Cell Culture Flask with a solid cap. The flasks were filled to the brim, leaving only a small headspace volume. The cells were concentrated by gently agitating the flask sediment, without disturbing the amoebae, and decanting 90% of the culture volume. The cells were dislodged using a cell scraper, and the concentrated cell solution was applied in a shallow glass beaker containing several Sapphire discs. The glass beaker itself was set in a sterile glass jar. The cells were allowed to sediment and attach for 10 min, after which culture supernatant from the same 75 cm^2^ culture flask was reapplied to fill the glass jar to the top. The cells were incubated for a minimum of 1 hour, and light microscopy of the immersed Sapphire disc showed high local amoebae density with spreading cells on the Sapphire discs.

### FIB-SEM

Cells were grown on Sapphire discs and high-pressure frozen in 100 µm deep golden carriers in an HPM100 (Leica microsystems, Wetzlar, Germany) with 20% BSA in 1xASW as cryo-protectant. Freeze substitution was performed as in Guo et al^16,^ but with shorter washing steps for the sake of time. After freeze substitution, the samples were infiltrated with TAAB embedding resin hard grade (TAAB Laboratories, Aldermaston, England). The sample was mounted on an SEM-stub with epoxy and silver glue. The sample was further coated with 5 nm Pt to reduce charging. Volumes were acquired using a Scios dualbeam (FEI, Eindhoven, The Netherlands) with the electron beam operating at 2 kV/0.2 nA, detected with the T1 In-lens detector. To automate the acquisition, we use the Auto Slice and View 4 software provided with the microscope. A 700 nm protective layer of platinum was deposited on the selected area before milling. The volume was further registered and processed by the ImageJ plugins Linear alignment by SIFT and Multistackreg. After registration, the volumes were converted to mrc-files, and the header was modified to recover the pixel size that got lost during conversion. The resolution of the two volumes and the number of slices were: 9235_1 (8.43 nm x 8.43 nm x 8.0 nm), 1,725 slices, and 9235_2 (6.744 nm x 6.744 nm x 7.0 nm), 1,300 slices.

### Segmentation

The *A. flamelloides* BUSSELTON2 cells were segmented by a combination of manual segmentation and Deep Learning Segmentation in Microscopy Image Browser v2.91 beta 45^17,18^. Training sets were assembled using the Segment Anything Model 2 (SAM 2). The training dataset for *endoplasmic reticulum and endocytic compartments* was assembled by using the whole model and gradually using the different individual models to cut out the model. The final cutout was introduced to the mask and used for mask-restricted BW thresholding to yield the respective training slices. Briefly, *Desulfobacter sp. symbionts, whole cells, symbiosome, endoplasmic reticulum and endocytic compartments, and hydrogenosomes* were manually annotated throughout the volume every 50-slices where the FIB-SEM volume was abundant in those structures. Image segments and the central model (patch size 256×256) were extracted 5 slices deep and used to train using the 2.5D semantic segmentation approach using 5 slices deep in Z2C+DLv3 architecture with the ResNet50 model. The predicted structures were manually refined extensively. *Nucleus* – segmented using the SAM 2 model, interactive 3D model. *Nuclear pores* – The nucleus model was dilated by 15 pixels, and BW thresholding was used to select the pixels that corresponded to nuclear pores, and was manually refined. *Plasma membrane and symbiosome membrane*– the whole cell and symbiosome model, respectively, were used to create an eroded mask to create a 5 px cutout of the respective membranes. Sharply shifting membrane sections were manually refined. *Microtubule Organizing Centre and microtubules* – the MTOC was manually segmented using SAM 2. Microtubules were traced manually using a 3 px brush. *Other structures (digestive vacuoles/inclusions/secondary symbionts)* – manually segmented using the SAM 2 model. *Symbiosome-to-surface openings* – symbiosome openings were identified manually in all three orientations and labeled using 3D balls. Three model files (exported from MIB^17,18^) were compiled for each volume, one for the whole cell, one for the full symbiosome and one with the remaining segmentations (9235_1, 10 segments: nucleus, nuclear pores, endoplasmic reticulum and endocytic compartments, hydrogenosomes, symbiosome membrane, other structures (digestive vacuoles/inclusions/secondary symbionts), *Desulfobacter* sp. symbionts, plasma membrane, microtubule organizing centre and microtubules, symbiosome-to-surface openings and 9235_2, 7 segments: nucleus, nuclear pores, hydrogenosomes, symbiosome membrane, *Desulfobacter* sp. symbionts, plasma membrane, symbiosome-to-surface openings.) The segmentations were rendered in Dragonfly v.2022.2.0 Build 1399^19^.

## Data availability

Aligned FIB-SEM .mrc files and segmentation models in .model format are available from the SciLifeLab Data Repository (https://doi.org/10.17044/scilifelab.31894429).

## Acknowledgement

The author acknowledges Umeå Centre for Electron Microscopy (UCEM) for technical assistance and access to electron microscopy. Service/Support/Collaboration was provided/supported by SciLifeLab Integrated Microscopy Technology Unit at Umeå University.

## Figures and supplementary material

**Figure S1:**
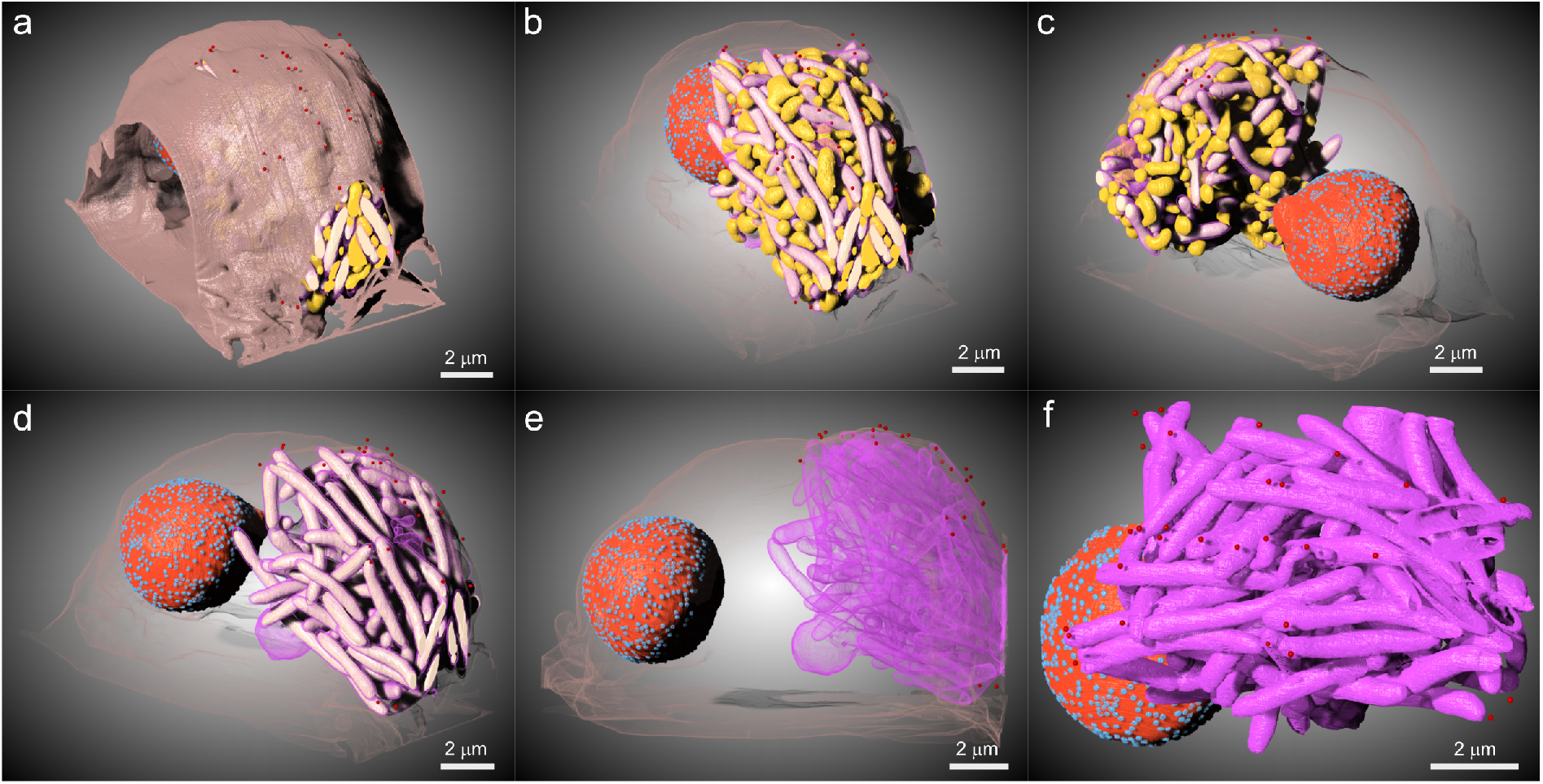
High-pressure frozen cell of *A. flamelloides* BUSSELTON2 with well-preserved membrane-structures. **a-f**, Renderings of selected segmented organelles/cell structures of *A. flamelloides* BUSSELTON2 volume 9235_2, showing (symbiont – white, hydrogenosome – yellow, symbiosome membrane – purple, nucleus – red, nuclear pores – turquoise, plasma membrane – light brown, symbiosome-to-surface openings – dark red). **a**, symbiont, hydrogenosome, symbiosome membrane, nucleus, nuclear pores, plasma membrane, symbiosome-to-surface openings. **b**, as in **a**, but with plasma membrane shaded. **c**, same as in **b**, but the cell is facing sideways and zoomed in. **d**, same as in **c**, but without hydrogenosomes. **e**, symbiosome, nucleus, nuclear pores, and plasma membrane shaded and symbiosome-to-surface openings with the cell facing sideways. **f**, symbiosome, nucleus, nuclear pores, and symbiosome-to-surface openings. Scale bar 2 µm.

**Figure S2:**
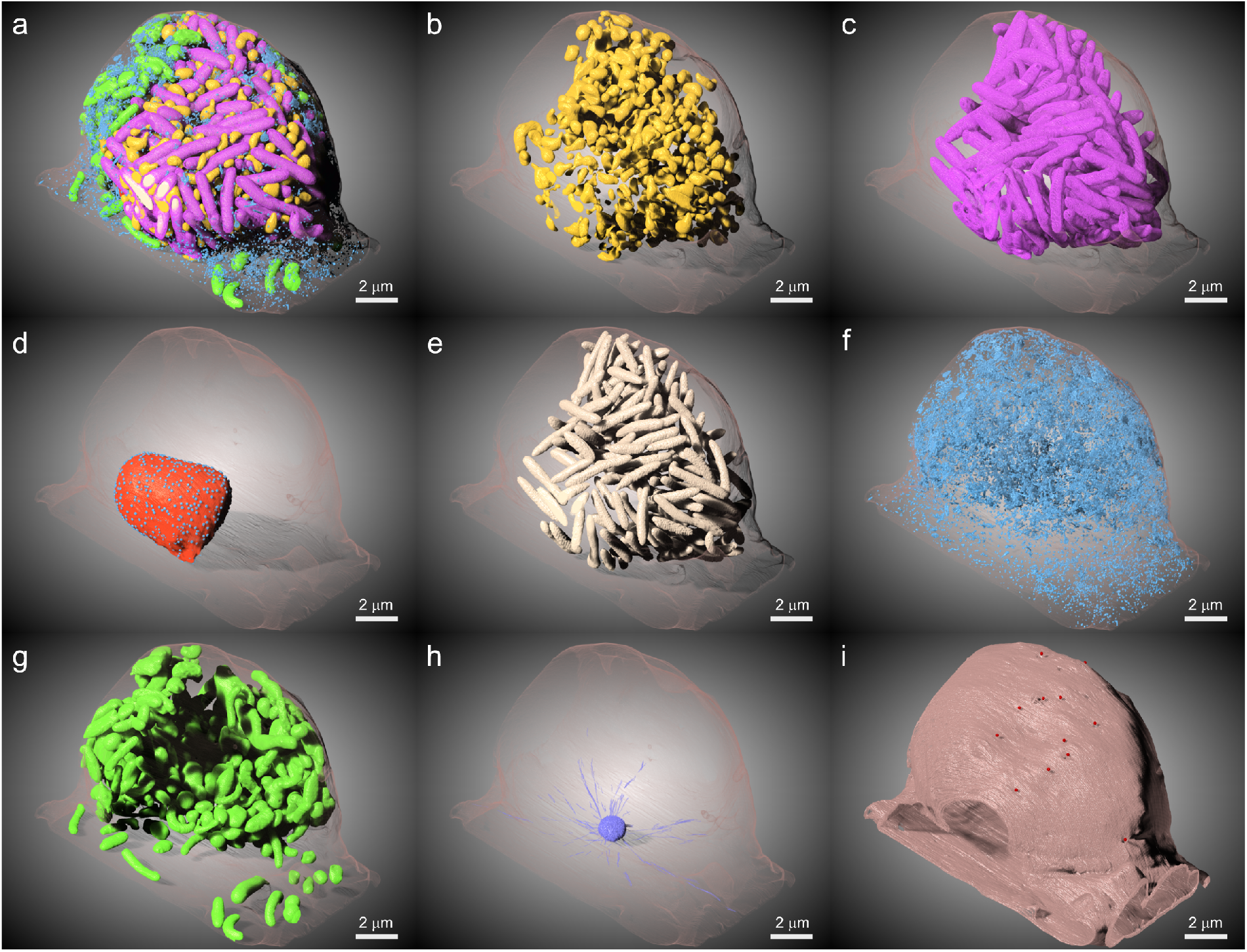
All segments of *A. flamelloides* BUSSELTON2 cell 9235_1. **a-i**, Renderings of organelles/cell structures of *A. flamelloides* BUSSELTON2 volume 9235_1, showing (symbiont – white, hydrogenosome – yellow, symbiosome membrane – purple, nucleus – red, nuclear pores – turquoise, plasma membrane – light brown, symbiosome-to-surface openings – dark red, MTOC and microtubules – dark blue, endoplasmic reticulum and endocytic compartments – blue, other structures (vacuoles, bacteria, inclusions - green). **a**, symbiont, hydrogenosome, symbiosome membrane, nucleus, nuclear pores, plasma membrane, symbiosome-to-surface openings, MTOC, other structures, endomembrane system, plasma membrane (shaded). **b-h**, plasma membrane is shaded and shown with **b**, hydrogenosomes, **c**, symbiosome, **d**, nucleus and nuclear pores, **e**, symbionts, **f**, endoplasmic reticulum and endocytic compartments, **g**, other structures, **h**, MTOC and microtubules. **i**, plasma membrane, and symbiosome-to-surface openings. Scale bar 2 µm.

**Figure S3:**
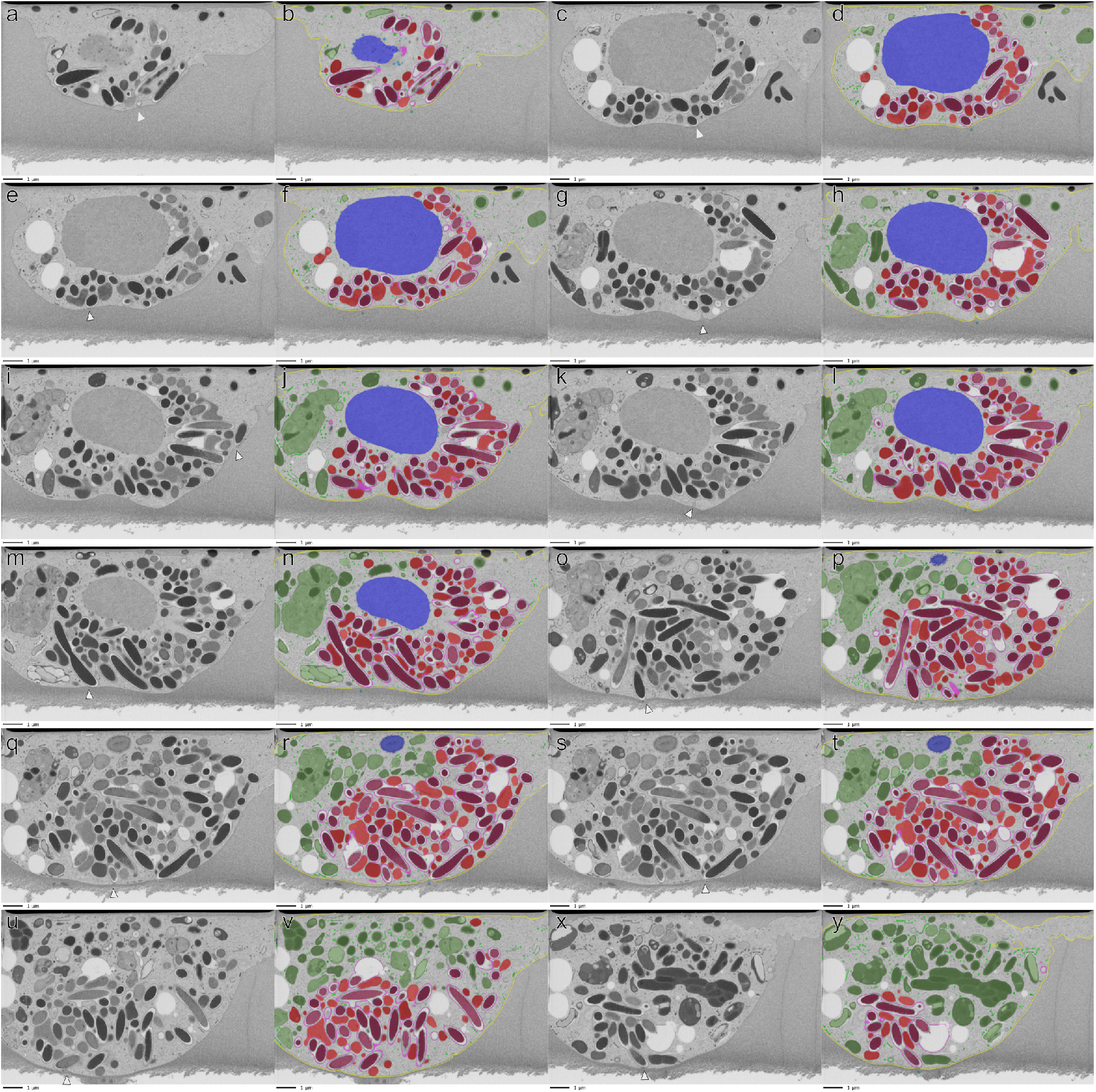
Symbiosome-to-surface openings in *A. flamelloides* BUSSELTON2 volume 9235_1. Paired FIB-SEM slices showing slices with symbiosome-to-surface openings indicated. Raw FIB-SEM slices in **a, c, e, g, i, k, m, o, q, s, u, x**. Symbiosome-to-surface openings are indicated by a white arrowhead. FIB-SEM slices with segmented structures in **b, d, f, h, j, l, n, p, r, t, v, y**. (*Desulfobacter* sp. symbiont – mauve, hydrogenosome – red, symbiosome membrane – purple, nucleus – blue, nuclear pores - light blue, other structures – dark green, plasma membrane – yellow, endoplasmic reticulum and endocytic compartments – light green, microtubule organizing centre – dark blue, symbiosome-to-surface openings - blue green dots). Scale bar 1 µm.

**Table S1:** Segmentation statistics of *A. flamelloides* BUSSELTON2; 9235_1 and 9235_2 volumes. The number of organelles, labeled voxels, and volume (cell%). The voxel statistics were calculated by Dragonfly v.2022.2.0 Build 1399^19^.

| **Segmentations of volume 9235_1** | **Organelles** | **Labeled voxels** | **Volume (cell %)** |
| --- | --- | --- | --- |
| Whole cell |  | 1623341799 | 100,0000 |
| Symbiosome full |  | 248601414 | 15,3142 |
| 1. Nucleus | 1 | 90599858 | 5,5811 |
| 2. Nuclear pores |  | 626966 | 0,0386 |
| 3. Endoplasmic reticulum and endocytic compartments |  | 18882169 | 1,1632 |
| 4. Hydrogenosomes | 374 | 102455814 | 6,3114 |
| 5. Symbiosome membrane | 1 | 37717471 | 2,3234 |
| 6. Other structures | N/A | 264425669 | 16,2890 |
| 7. *Desulfobacter* sp. symbionts | 160 | 112891533 | 6,9543 |
| 8. Plasma membrane | 1 | 35945209 | 2,2143 |
| 9. MTOCe and microtubules | N/A | 1388012 | 0,09 |
| 10. Symbiosome-to-surface openings | 12 |  |  |
| **Segmentations of volume 9235_2** | **Number of organelles** | **Labeled voxels** | **Volume (cell %)** |
| Whole cell |  | 2937263040 | 100,0000 |
| Symbiosome full |  | 325559232 | 11,0838 |
| 1. Nucleus | 1 | 191921097 | 6,5340 |
| 2. Nuclear pores |  | 1306602 | 0,0445 |
| 3. Endoplasmic reticulum and endocytic compartments |  | not segmented |  |
| 4. Hydrogenosomes | 307 | 179041792 | 6,0955 |
| 5. Symbiosome membrane |  | 47310218 | 1,6107 |
| 6. Other structures | N/A | not segmented |  |
| 7. *Desulfobacter* sp. symbionts | 95 | 150153337 | 5,1120 |
| 8. Plasma membrane |  | 49607229 | 1,6889 |
| 9. MTOC and microtubules | N/A | not segmented |  |
| 10. Symbiosome-to-surface openings | 29 |  |  |

